# FRA1 modulates cortical microtubule localization of CMU proteins

**DOI:** 10.1101/760389

**Authors:** Anindya Ganguly, Chuanmei Zhu, Weizu Chen, Ram Dixit

## Abstract

Construction of the cell wall demands harmonized deposition of cellulose and matrix polysaccharides. Cortical microtubules orient the deposition of cellulose by guiding the trajectory of plasma membrane-embedded cellulose synthase complexes. Vesicles containing matrix polysaccharides are thought to be transported by the FRA1 kinesin to facilitate their secretion along cortical microtubules. The cortical microtubule cytoskeleton thus provides a platform to coordinate the delivery of cellulose and matrix polysaccharides, but the underlying molecular mechanisms remain unknown. Here, we show that the tail region of the FRA1 kinesin physically interacts with CMU proteins which are important for the microtubule-dependent guidance of cellulose synthase complexes. Interaction with CMUs did not affect microtubule binding or motility of the FRA1 kinesin but had an opposing effect on the cortical microtubule localization of CMU1 and CMU2 proteins, thus regulating the lateral stability of cortical microtubules. Phosphorylation of the FRA1 tail region by CKL6 inhibited binding to CMUs and consequently reversed the extent of cortical microtubule decoration by CMU1 and CMU2. Genetic experiments demonstrated the significance of this interaction to the growth and reproduction of *Arabidopsis thaliana* plants. We propose that modulation of CMU’s microtubule localization by FRA1 provides a mechanism to control the coordinated deposition of cellulose and matrix polysaccharides.

## INTRODUCTION

Plants build a mechanically tough cell wall which confers cell shape by defining the direction of turgor-driven expansion. The cell wall is a composite material consisting primarily of cellulose microfibrils enmeshed within a complex network of hemicellulose and pectin. Cells control the supply locations and delivery patterns of these components to regulate wall mechanics and thus spatially guide their growth. A key component of this control mechanism is the cortical microtubule cytoskeleton, which serves as a scaffolding structure that spatially organizes the cell wall synthesis machinery. Disruption of the cortical microtubule array by genetic or pharmacological methods perturbs wall assembly and leads to aberrant plant growth and development.

Cellulose microfibrils are synthesized at the cell surface by transmembrane cellulose synthase complexes (CSCs). CSCs contained in Golgi-derived membrane compartments are inserted into the plasma membrane adjacent to cortical microtubules (Crowell et al., 2009; Gutierrez et al., 2009). Catalytically active CSCs move steadily along cortical microtubules (Paredez et al., 2006) due to physical linkage provided by CELLULOSE SYNTHASE INTERACTIVE proteins (Bringmann et al., 2012; Li et al., 2012). The cortical microtubule associated CELLULOSE SYNTHASE-MICROTUBULE UNCOUPLING proteins (CMUs) in turn maintain the positional stability of cortical microtubules in the face of forces generated by motile CSCs (Liu et al., 2016). Together, these mechanisms constitute a multi-step process by which cortical microtubules orient cellulose deposition.

Hemicellulose and pectin are synthesized in the Golgi and secreted into the extracellular space for incorporation in the cell wall. Accumulating evidence indicates that deposition of at least some of these matrix polysaccharides is directed by cortical microtubules (Fukuda, 1997; McFarlane et al., 2008; Kong et al., 2015; Zhu et al., 2015; Takenaka et al., 2018). Recent work on the *Arabidopsis thaliana* FRAGILE FIBER1 (FRA1) kinesin has uncovered a potential molecular mechanism for this process. Kinesins are ATP-dependent molecular motors that transport cellular cargo along microtubules in a directional manner. The FRA1 kinesin was found to move long distances along cortical microtubules (Zhu and Dixit, 2011; Kong et al., 2015; Zhu et al., 2015) and this property was shown to be essential for its function (Ganguly et al., 2017). Loss of the FRA1 kinesin leads to accumulation of vesicles in the vicinity of Golgi bodies and greatly reduced pectin secretion (Zhu et al., 2015). Based on these data, the FRA1 kinesin is thought to transport secretory vesicles containing matrix polysaccharides to exocytic sites positioned along cortical microtubules.

The deposition of cellulose and matrix polysaccharides needs to be coordinated spatially and temporally for proper cell wall assembly. When the demand for cell wall synthesis rises, such as during rapid cell elongation, the abundance and motility of both CSCs and the FRA1 kinesin increases proportionally (Crowell et al., 2009; Zhu et al., 2015; Ganguly et al., 2018). However, the mechanisms for the concurrent regulation of cellulose and matrix polysaccharide deposition remain unknown.

Here, we report that CMUs directly interact with the tail region of the FRA1 kinesin and together they contribute to plant stature and fertility. We find that the cortical microtubule localization of CMUs is regulated by their interaction with FRA1, whereas the motility and function of the FRA1 kinesin is unaffected by CMUs. Finally, we show that phosphorylation of the FRA1 tail domain by CASEIN KINASE-LIKE6 (CKL6) weakens its interaction with CMUs, providing a mechanism to tune the extent of cortical microtubule localization of CMUs. Together, our data reveal a new mechanism that functionally links the cellulose and matrix polysaccharide deposition machinery.

## RESULTS

### The FRA1 kinesin physically interacts with CMU proteins

A yeast two-hybrid screen conducted by Hybrigenics Services with a C-terminal fragment of the FRA1 kinesin that includes the tail domain and part of the stalk region (amino acids 616-1035) (Figure 1A) identified the *Arabidopsis thaliana* CMU1 and CMU2 proteins as putative interacting partners. Based on amino-acid sequence similarity to the human kinesin light-chain proteins, the Arabidopsis CMUs were also previously called kinesin light chain-related proteins (Burstenbinder et al., 2013). We confirmed interaction between the FRA1 C-terminal fragment and full-length CMU1 and CMU2 proteins using directed yeast two-hybrid assays (Figure 1B). In addition, we performed *in vitro* pull-down experiments and found that all three Arabidopsis CMU proteins interact with the FRA1 C-terminal fragment (Figure 1C and Supplemental Figure 1A). To determine whether this interaction occurs *in vivo*, we immunoprecipitated mCherry-tagged CMU1 and CMU2 expressed in wild-type *A. thaliana* plants under their respective native promoters. We found that both mCherry-CMU1 and mCherry-CMU2 interact with full-length FRA1 in plants (Figure 1D). Similar experiments with mCherry-CMU3 were unsuccessful, probably because CMU3 expression is below the detection level in immunoblots (Supplemental Figure 1B). Together, these results identify CMUs as authentic interaction partners of the FRA1 kinesin.

**Figure 1.**
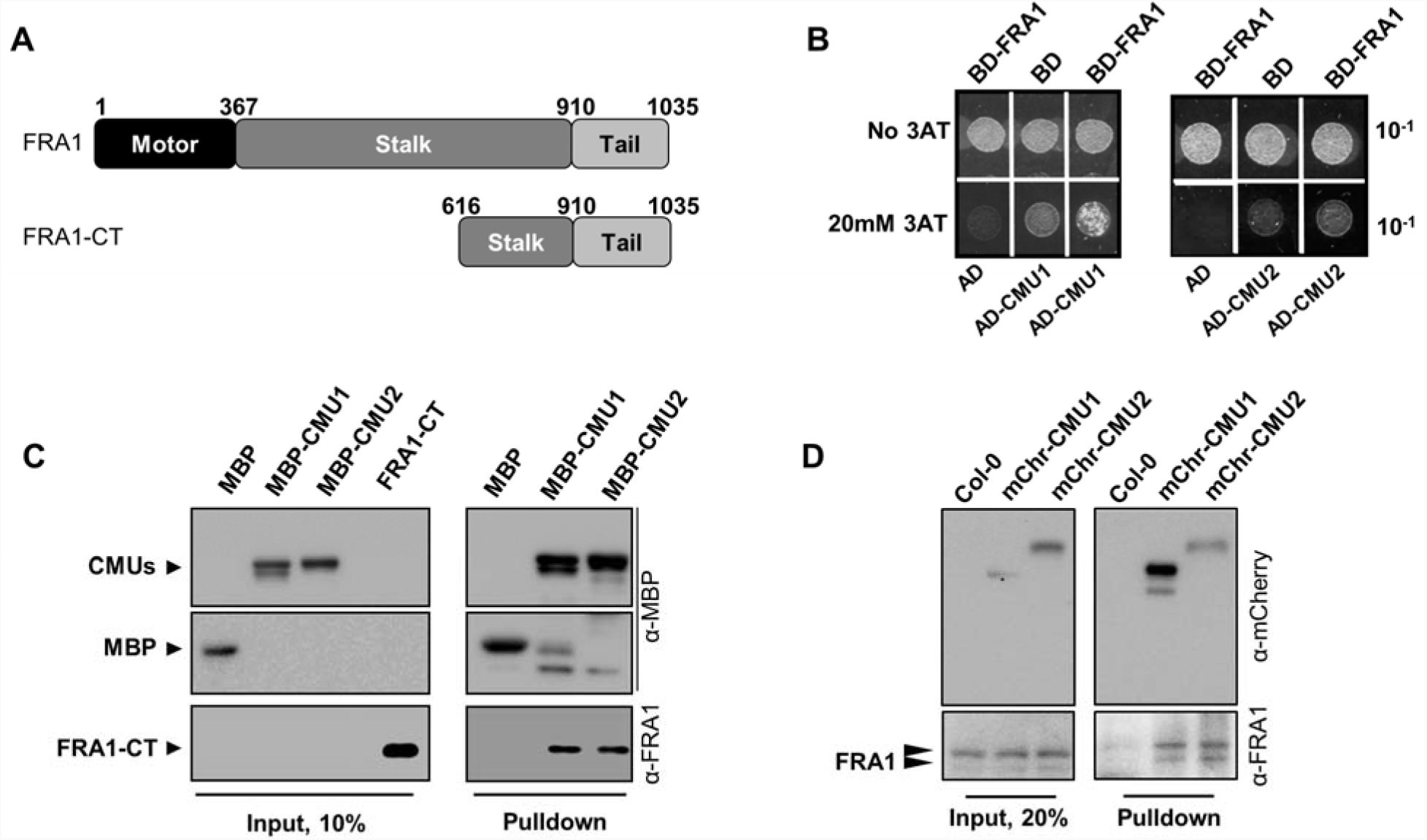
FRA1 interacts directly with CMU1 and CMU2. (A) Schematic representation of the FRA1 full-length protein and C-terminal region (FRA1-CT) used for pulldown and yeast two-hybrid assays in this study. (B) Yeast two-hybrid assay using FRA1-CT fused with the activation domain (AD) as prey and CMU1 and CMU2 fused with the DNA binding domain (BD) as bait. Cells were grown on plates without or with 20 mM 3-amino-1,2,4-triazole (AT). (C) Pulldown of either MBP alone or MBP-tagged CMU1 and CMU2 proteins incubated with equal amounts of FRA1-CT. Immunoblotting was performed with anti-MBP and anti-FRA1 antibodies. (D) Pulldown of mCherry-tagged CMU1 (mChr-CMU1) and CMU2 (mChr-CMU2) proteins expressed in Col-0 plants using their native promoters. Immunoblotting was performed with anti-mCherry and anti-FRA1 antibodies. The endogenous FRA1 protein frequently shows up as a doublet in immunoblots, as previously reported (Zhu et al., 2015 and Ganguly et al., 2018).

### Loss-of-function *cmu* mutants enhance the growth and fertility defects of the *fra1-5* mutant

Previous work had identified transfer-DNA (T-DNA)-induced loss-of-function mutants of *CMU1* and *CMU2*, named *cmu1* and *cmu2* (Liu et al., 2016). Since T-DNA insertion alleles are unavailable for *CMU3*, we used artificial microRNA technology to knockdown *CMU3* expression in the *cmu1cmu2* mutant background (Supplemental Figure 1C).

We found that *cmu1* seedlings have skewed roots whereas *cmu2* seedlings are indistinguishable from wild-type (Supplemental Figure 1D-1E). The *cmu1cmu2* double mutant and *cmu1cmu2cmu3* triple mutant have skewed roots similar to the *cmu1* single mutant (Supplemental Figure 1D-1E). These data indicate that CMU1 prevents root skewing in seedlings whereas CMU2 and CMU3 do not contribute to this function. In the adult plant stage, none of the *cmu* mutants showed significant growth defects compared to wild type plants (Figure 2A-2B and Supplemental Figure 1F-1G).

**Figure 2.**
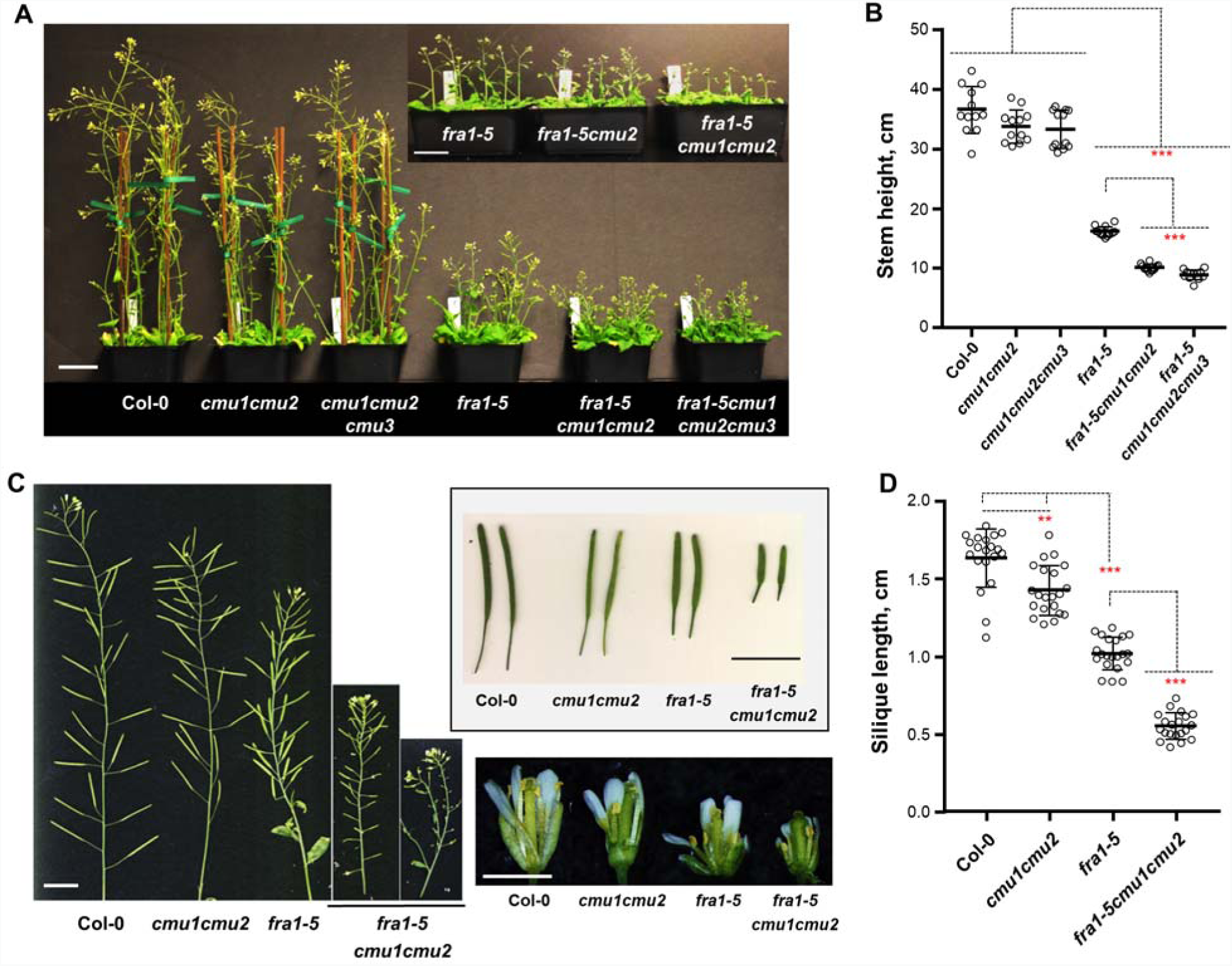
Phenotypes of *cmu* and *fra1-5* mutant combinations. (A, B) Whole-plant appearance and quantification of stem heights of plants 42 days after germination. Bars indicate mean ± S.D. (n > 12 plants). Scale bar = 5 cm. (C) Images of primary inflorescence stems, isolated siliques and flowers. Scale bar = 1 cm. (D) Scatterplot of silique length. Bars indicate mean ± S.D. (n > 20 mature siliques). Asterisks indicate significant difference between the indicated genotypes as determined by ANOVA, **p < 0.001, ***p < 0.0001.

To determine whether the physical interaction between FRA1 and CMUs is functionally important, we crossed the *cmu* mutants with the previously described *fra1-5* knockout mutant (Zhu et al., 2015). We found that the root skewing phenotype of the *cmu1* single mutant was partially suppressed in the *fra1-5cmu1* double mutant (Supplemental Figure 1D-1E). However, the dwarf phenotype of the *fra1-5cmu1* double mutant was indistinguishable from the *fra1-5* single mutant (Supplemental Figure 1F-1G). In contrast, the *fra1-5cmu2* double mutant was significantly shorter than the *fra1-5* mutant (Supplemental Figure 1F-1G). Stem height was further reduced in the *fra1-5cmu1cmu2* triple mutant (Figure 2A-2B), indicating that CMU1 and CMU2 contribute additively to stem height in the *fra1-5* background. In contrast, knockdown of *CMU3* did not further decrease stem height in the *fra1-5cmu1cmu2cmu3* quadruple mutant (Figure 2A-2B).

While fertility of the *cmu1cmu2* double mutant is similar to wild-type plants, we found that it greatly enhanced the fertility defect of the *fra1-5* mutant. As previously reported, the *fra1-5* mutant has ~10% undeveloped siliques compared to ~1% undeveloped siliques in wild-type plants (Kong et al., 2015). Introgression of the *cmu1cmu2* double mutation into the *fra1-5* line resulted in ~30% undeveloped siliques in the *fra1-5cmu1cmu2* triple mutant (Figure 2C). In addition, fully developed siliques of the *fra1-5cmu1cmu2* triple mutant were about half the length of the *fra1-5* siliques (Figure 2D). To determine whether paternal or maternal tissues contribute to reduced fertility of these mutants, we performed reciprocal pollination experiments and measured the extent of pollen tube growth in the transmitting tract of the pistil. When wild-type pistils were used, we found that pollen grains from the *cmu1cmu2* double mutant grew to a similar length as wild-type pollen (Figure 3A-3B). In contrast, pollen grains from the *fra1-5* single mutant and the *fra1-5cmu1cmu2* triple mutant grew significantly less over the same time period (Figure 3A-3B). In the reciprocal experiments, wild-type pollen grew to a lesser extent on *fra1-5* pistils compared to on wild-type and *cmu1cmu2* pistils and this defect was exacerbated on *fra1-5cmu1cmu2* pistils (Figure 3C-3D).

**Figure 3.**
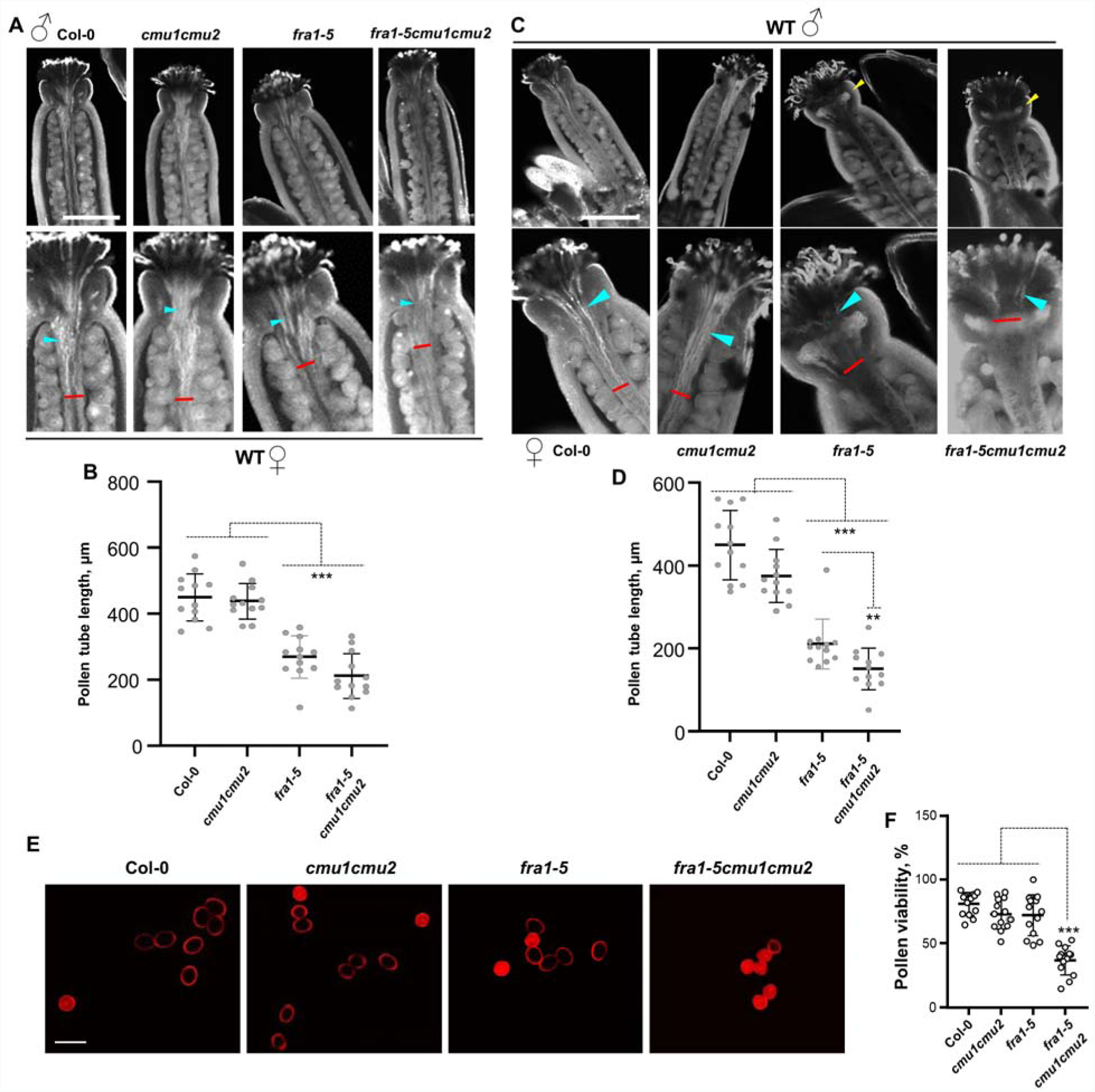
Pistil and pollen defects in *cmu* and *fra1-5* mutant combinations. (A) Images of aniline blue-stained wild-type pistils after 2-h pollination with pollen from the indicated genotypes. Magnified views are shown in the bottom panel. Blue arrowheads point to pollen tubes growing in the transmitting tract. Red bars indicate the distance of pollen tube growth. Scale bar = 250 µm. (B) Scatterplot of pollen tube lengths from experiments shown in (A). Bars indicate mean ± S.D. (n > 12 sets). (C) Images of aniline blue-stained pistils of the indicated genotype after 2-h pollination with wild-type pollen. Magnified views are shown in the bottom panel. Blue arrowheads point to pollen tubes growing in the transmitting tract. Red bars indicate the distance of pollen tube growth. Pistils of the *fra1-5* mutant reproducibly show a band of callose beneath the stigma (yellow arrowhead). Scale bar = 250 µm. (D) Scatterplot of pollen tube lengths from experiments shown in (C). Bars indicate mean ± S.D. (n > 12 sets). (E) Micrographs of pollen grains stained with 1 µg/ml propidium iodide for 15 min. The dye is excluded from the cytoplasm of viable pollen. Scale bar = 20 µm. (F) Scatterplot of the percentage of viable pollen grains. Bars indicate mean ± S.D. (n > 12 sets with at least 15 pollen in each set) Asterisks indicate significant difference between the indicated genotypes as determined by ANOVA, **p < 0.001, ***p < 0.0001.

During these experiments, we observed that pollen grains from the *fra1-5cmu1cmu2* triple mutant often failed to germinate. To determine whether this was due to reduced pollen viability, we stained pollen grains with propidium iodide. We found that while pollen grains from the *fra1-5* and *cmu1cmu2* mutants showed similar viability as wild-type, pollen grains from the *fra1-5cmu1cmu2* triple mutant showed ~ 30% decrease in viability (Figure 3E-3F). Together, our data indicate that the CMU1 and CMU2 proteins work together with the FRA1 kinesin to achieve normal pollen and pistil function.

### Motility of the FRA1 kinesin remains unaffected in the *cmu1cmu2* mutant

In mammals, kinesin light chains regulate the processive motility of kinesin-1 motors (Verhey et al., 1998; Cai et al., 2007; Wong and Rice, 2010). Since CMUs resemble kinesin light chains, we investigated whether the motility of FRA1 is altered in the *cmu1cmu2* double mutant. For this purpose, we generated a single-GFP tagged version of full-length FRA1 driven by its own promoter and introduced it in the *fra1-5* and *fra1-5cmu1cmu2* mutants. The *pFRA1::FRA1-GFP* construct fully complemented the dwarf phenotypes of both of these mutants, indicating that it is functional (Figure 4A-4B). Live imaging of hypocotyl epidermal cells revealed that FRA1-GFP particles move processively in both *fra1-5* and *fra1-5cmu1cmu2* mutants (Figure 4C). We generated kymographs to quantify motility of the FRA1-GFP particles and found that their velocity, run length and motile density were indistinguishable in the *fra1-5* and *fra1-5cmu1cmu2* backgrounds (Figure 4D-4E). Therefore, we conclude that CMU1 and CMU2 do not regulate motility of the FRA1 kinesin.

**Figure 4.**
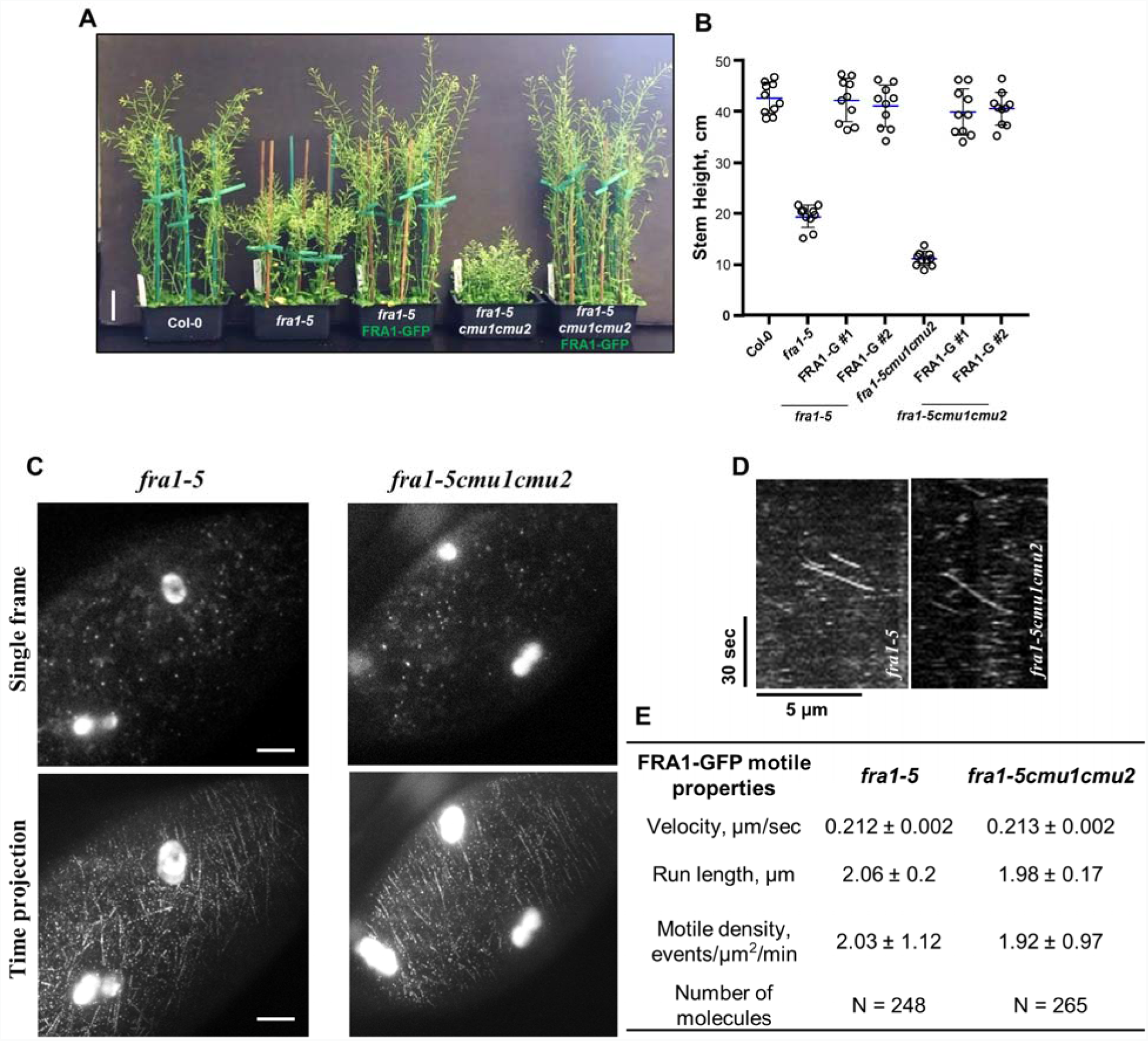
CMUs do not regulate the motility of FRA1. (A, B) Whole-plant appearance and quantification of stem heights of plants 42 days after germination. Bars indicate mean ± S.D. (n = 12 plants each). Scale bar = 5 cm. (C) Single frames and time projections of 120 frames of FRA1-GFP. Directional movement of FRA1-GFP molecules appear as linear tracks in the time projection. The bright oval structures in these images are chloroplasts imaged because of chlorophyll autofluorescence. Scale bar = 5 µm. (D) Representative kymographs showing the movement of FRA1-GFP. Diagonal lines represent motile events. (E) Motile parameters of processive FRA1-GFP puncta. Values are mean ± S.D.

### FRA1 differentially regulates CMU1 and CMU2 cortical microtubule localization

To determine if FRA1 affects CMU proteins, we generated constructs that express mCherry-tagged CMU1 and CMU2 under the control of their respective native promoters. We found that mCherry-CMU1 and mCherry-CMU2 fusion proteins colocalize with cortical microtubules (Supplemental Figure 2A), as reported previously (Liu et al., 2016). Importantly, both fusion proteins are functional because they fully complement the growth defects of *cmu1* and *fra1-5cmu2* mutants, respectively (Supplemental Figure 2B-2E). To examine the effect of FRA1 on these proteins, we introduced the mCherry-CMU1 and mCherry-CMU2 markers into the *fra1-5* mutant. We found that the signal of mCherry-CMU1 along cortical microtubules was significantly reduced in the *fra1-5* mutant compared to wild-type (Figure 5A-5B). Conversely, the signal of mCherry-CMU2 along cortical microtubules was significantly enhanced in the *fra1-5* mutant compared to wild-type (Figure 5C-5D). Complementation of the *fra1-5* mutant by *FRA1-GFP* restored the cortical microtubule signal of mCherry-CMU1 and mCherry-CMU2 to wild-type levels (Figure 5A and 5C).

**Figure 5.**
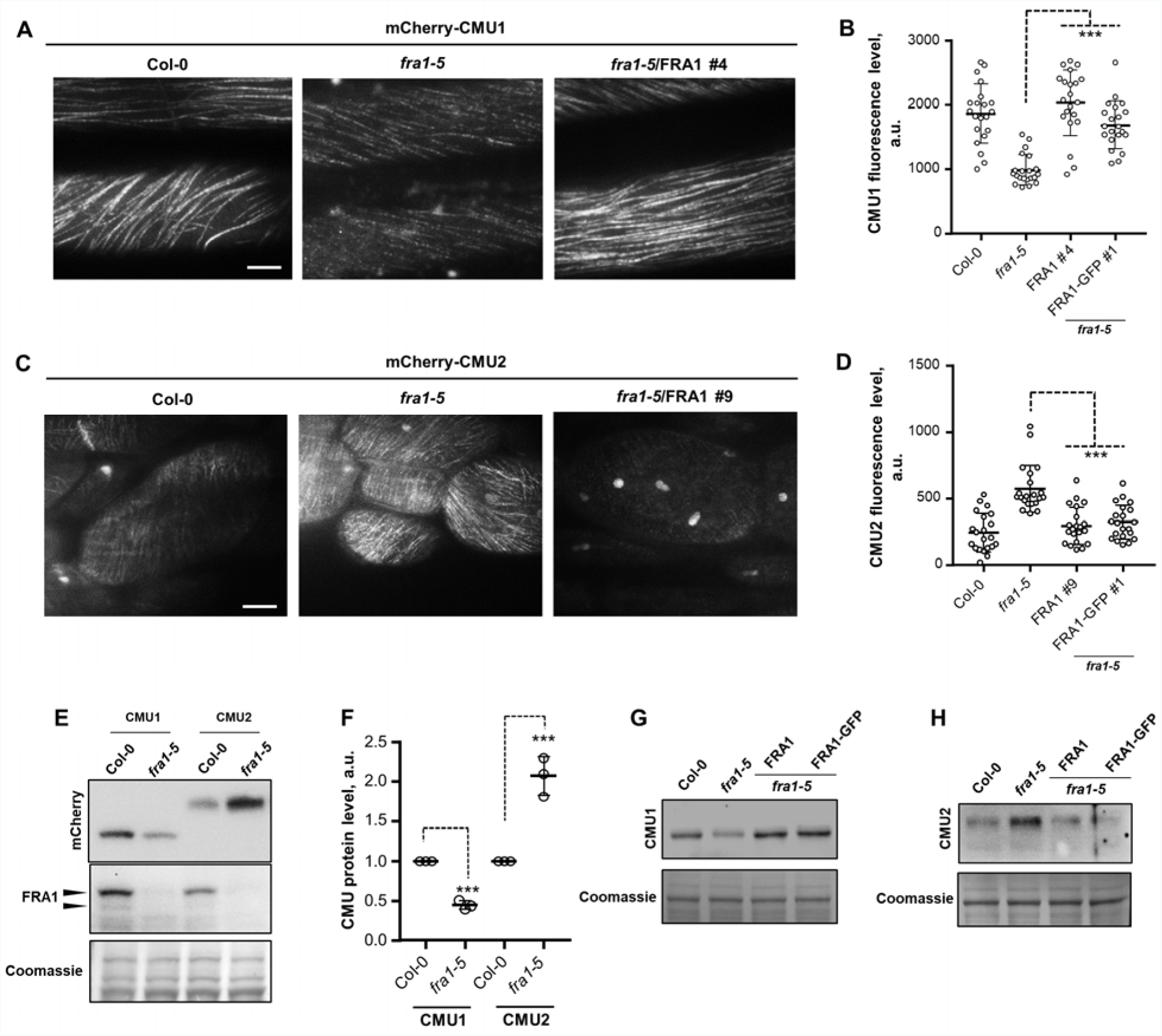
FRA1 differentially regulates CMU1 and CMU2 localization on cortical microtubules. All data are from 4-day-old seedlings. The immunoblots are representative of at least 3 independent experiments. Coomassie-stained gels are shown below each immunoblot as a loading control. (A-D) Fluorescence micrographs (A and C) and quantification (B and D) of mCherry-CMU1 and mCherry-CMU2 signal on cortical microtubules in hypocotyl epidermal cells. Values are mean ± S.D. (3-4 microtubules were quantified per cell, total 8 cells from 3 seedlings). Scale bar = 10 µm for (A) and 5 µm for (C). (E, F) Immunoblotting and quantification of CMU1 and CMU2 proteins levels in Col-0 and *fra1-5* plants. Total protein extracts were probed with anti-mCherry and anti-FRA1 antibodies. (G, H) Immunoblots of total protein extracts prepared from Col-0, *fra1-5* and *fra1-5* plants complemented with untagged FRA1 or FRA1-GFP probed with anti-mCherry antibody. Asterisks indicate significant difference between the indicated genotypes as determined by ANOVA, ***p < 0.0001.

To determine whether the changes in signal intensity of mCherry-CMU1 and mCherry-CMU2 are due to alteration of protein levels, we performed immunoblot analysis with an anti-mCherry antibody. These experiments revealed that the amount of mCherry-CMU1 protein was about 2-fold lower whereas the amount of mCherry-CMU2 protein was about 2-fold higher in the *fra1-5* mutant compared to wild-type plants (Figure 5E-5F). Consistent with our microscopy data, mCherry-CMU1 and mCherry-CMU2 protein levels returned to wild-type levels upon introduction of a functional copy of *FRA1* in the *fra1-5* mutant (Figure 5G-5H). Together, these data indicate that the FRA1 kinesin regulates the amount of CMU1 and CMU2 proteins available to bind to cortical microtubules.

### Lateral stability of cortical microtubules is reduced in the *fra1-5* mutant

CMU proteins prevent the lateral displacement of cortical microtubules by active CSCs (Liu et al., 2016). Since cortical microtubule localization of CMUs is altered in the *fra1-5* mutant, we wondered whether the lateral stability of cortical microtubules was compromised in *fra1-5* cells. Live imaging revealed that cortical microtubules in the *fra1-5* mutant showed lateral deflections similar to that observed in the *cmu1cmu2* double mutant (Figure 6A-6B). To measure the extent of lateral displacement of cortical microtubules, we used kymograph analysis as described previously (Liu et al., 2016). Cortical microtubules that stably maintain their position appear as vertical lines in kymographs, whereas cortical microtubules that drift laterally appear as oblique lines in kymographs (Figure 6C). We found that the extent and frequency of microtubule displacements were comparable between the *fra1-5* and *cmu1cmu2* mutants (Figure 6D-6E). In addition, this defect was not enhanced in the *fra1-5cmu1cmu2* triple mutant compared to the *fra1-5* and *cmu1cmu2* mutants (Figure 6D-6E). Together, these data indicate that FRA1 itself does not contribute to the positional stability of cortical microtubules and that this phenotype is likely the indirect consequence of lower CMU1 protein levels in the *fra1-5* mutant.

**Figure 6.**
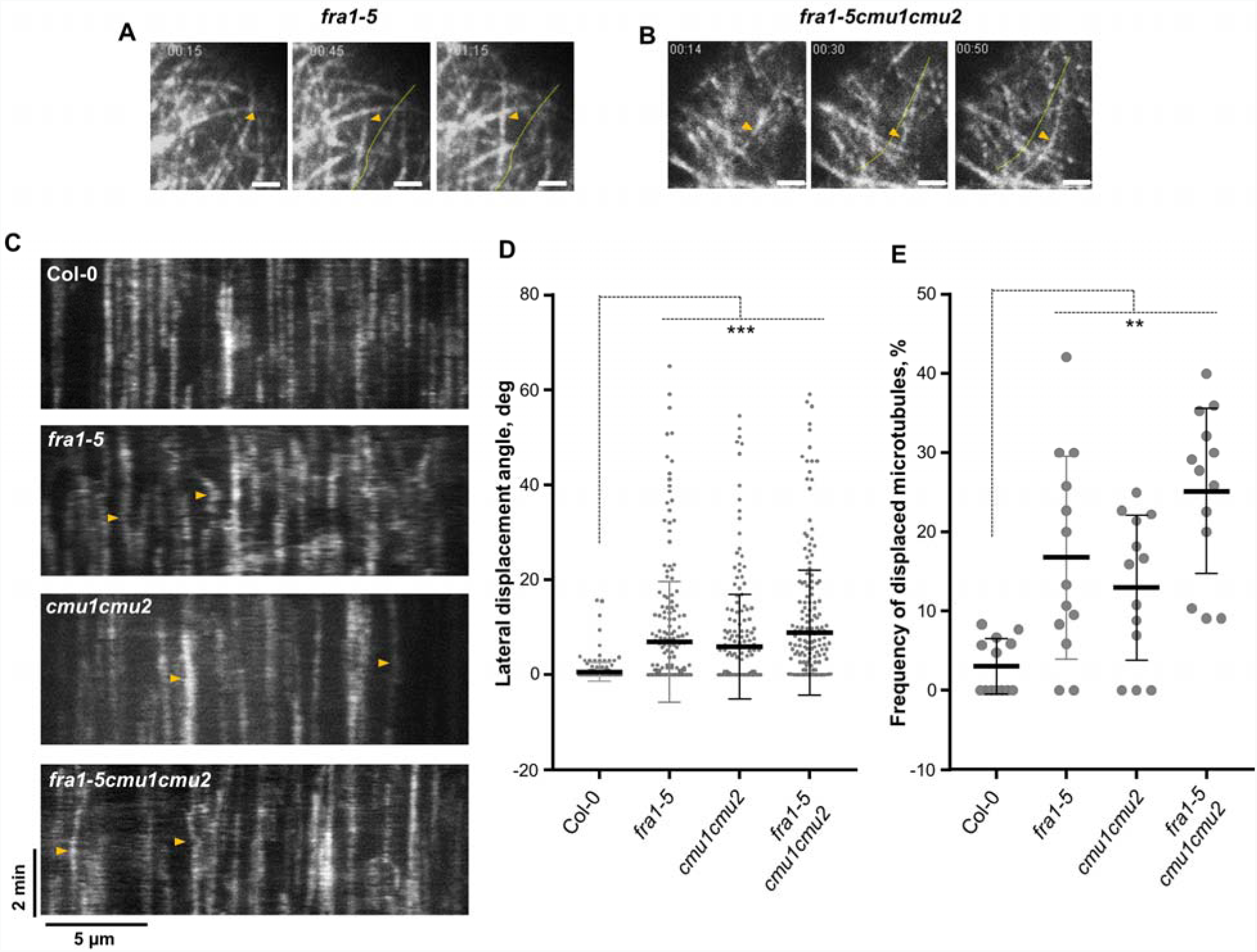
Lateral displacement of cortical microtubules in the *fra1-5* and *fra1-5cmu1cmu2* mutants. (A-B) Montage of single frames of cortical microtubules labeled with mRuby-TUB6 showing lateral displacement of a subset of cortical microtubules in the *fra1-5* and *fra1-5cmu1cmu2* mutants. The microtubule of interest is marked by an orange arrowhead. The dotted line in frames 2 and 3 show the starting position of the microtubule in frame 1. Scale bar = 2 µm. (C) Kymographs showing lateral displacement of cortical microtubules in hypocotyl epidermal cells. Positionally fixed microtubules appear as straight lines, whereas laterally drifting microtubules appear as curved lines (orange arrowheads). (D, E) Plots of the angle of lateral displacement of cortical microtubules (D) and the frequency of displaced microtubules (E) determined from kymographs. Bars indicate mean ± S.D. For D, n > 200 CMTs from 15-22 cells from 6 independent seedlings. For E, n = 12-13 cells from 5 independent seedlings). Asterisks indicate significant differences as determined by ANOVA, ** p < 0.001, *** p < 0.0001.

### The FRA1 C-terminal region is phosphorylated by casein kinase1-like 6

Mass spectrometry data in the Arabidopsis protein phosphorylation site database identified Thr687 and Thr694 to be phosphorylated in the FRA1 kinesin (Durek et al., 2010). These residues lie in the C-terminal region of FRA1 which interacts with CMUs and are predicted to be phosphorylated by kinases belonging to the casein kinase1-like family. To determine whether FRA1 is a substrate of these kinases, we selected casein kinase1-like 3 (CKL3) and casein kinase1-like 6 (CKL6) because they have been shown to be active kinases *in vitro* (Ben-Nissan et al., 2008; Tan et al., 2013). We performed *in vitro* pull-down experiments and found that full-length, MBP-tagged CKL3 and CKL6 both interact directly with the FRA1 C-terminal domain (Figure 7A). However, only CKL6 was able to phosphorylate the FRA1 C-terminal domain *in vitro* (Figure 7B). Mutating the Thr687 and Thr694 residues to alanine reduced phosphorylation of the FRA1 C-terminal domain by CKL6 by about 3-fold (Figure 7B). Control experiments showed that both CKL3 and CKL6 phosphorylate tubulin dimers *in vitro* (Supplemental Figure 3), demonstrating that the lack of phosphorylation of the FRA1 C-terminal domain by CKL3 was not due to lack of kinase activity.

**Figure 7.**
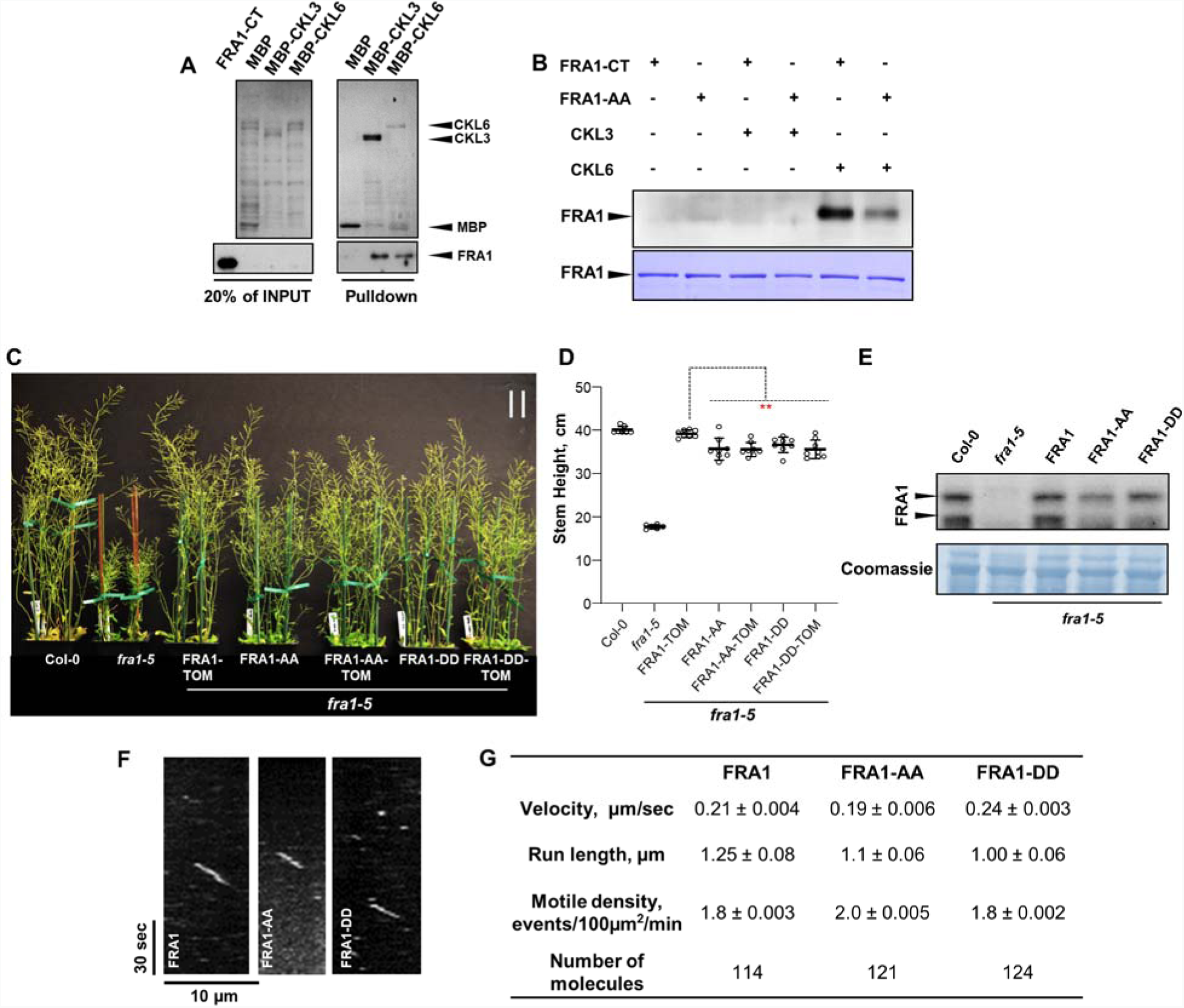
The C-terminal tail region of FRA1 is phosphorylated by CKL6. (A) Pulldown of either MBP alone or MBP-tagged CKL3 and CKL6 proteins incubated with equal amounts of FRA1-CT protein. Immunoblotting was performed with anti-MBP and anti-FRA1 antibodies. (B) *In vitro* phosphorylation experiment with wild-type FRA1-CT and FRA1-AA mutant incubated with either CKL3 or CKL6. Phosphorylation was detected by an anti-phospho serine/threonine antibody. Coomassie-stained gel is shown below as a loading control. (C, D) Whole-plant appearance and quantification of stem heights of plants 42 days after germination. Bars indicate mean ± S.D. (n = 12 plants each). Asterisks indicate significant difference as determined by ANOVA, **p < 0.005. Scale bar = 5 cm. (E) Immunoblot of total protein extracts probed with anti-FRA1 antibody. Coomassie-stained gel is shown below as a loading control. (F) Representative kymographs showing the movement of tdTomato-labeled wild-type FRA1, FRA1-AA and FRA1-DD mutants. Diagonal lines represent motile events. (G) Motile parameters of processive FRA1-tdTomato puncta. Values are mean ± S.D.

To study whether phosphorylation of FRA1 at these residues contributes to its function, we introduced untagged and tdTomato-tagged wild-type FRA1 or phosphodefective (FRA1-AA) or phosphomimetic (FRA1-DD) versions expressed under the control of the native *FRA1* promoter in the *fra1-5* mutant. In agreement with previous work (Ganguly et al., 2018), wild-type FRA1-tdTomato fully rescued the dwarf phenotype of the *fra1-5* mutant (Figure 7C-7D). In contrast, the phosphodefective and phosphomimetic FRA1 proteins restored stem height to ~ 90% of wild-type but did not completely rescue the *fra1-5* mutant (Figure 7C-7D). Importantly, the steady-state protein levels and motile properties of the FRA1 phosphorylation mutants were similar to wild-type FRA1 (Figures 7E-7G). Therefore, phosphorylation of Thr687 and Thr694 does not significantly affect the abundance and motility of the FRA1 kinesin.

### Phosphorylation of the FRA1 C-terminal region regulates the extent of CMU localization on cortical microtubules

Next, we investigated whether phosphorylation of Thr687 and Thr694 regulates the interaction of FRA1 with CMUs. *In vitro* pull-down experiments showed that the phosphodefective mutations of the FRA1 C-terminal region enhanced binding to CMU1 and CMU2 whereas the phosphomimetic mutations decreased binding to CMU1 and CMU2 (Figure 8A). To determine whether phosphorylation of FRA1 affects the microtubule localization of CMUs *in vivo*, we introduced untagged FRA1-AA and FRA1-DD proteins in the *fra1-5* mutant expressing either mCherry-CMU1 or mCherry-CMU2. We found that the phosphodefective FRA1 mutant restored the levels of mCherry-CMU1 and mCherry-CMU2 on cortical microtubules to wild-type levels (Figure 8B-8E). In contrast, the phosphomimetic FRA1 mutant was only partially able to perform this function (Figure 8B-8E).

**Figure 8.**
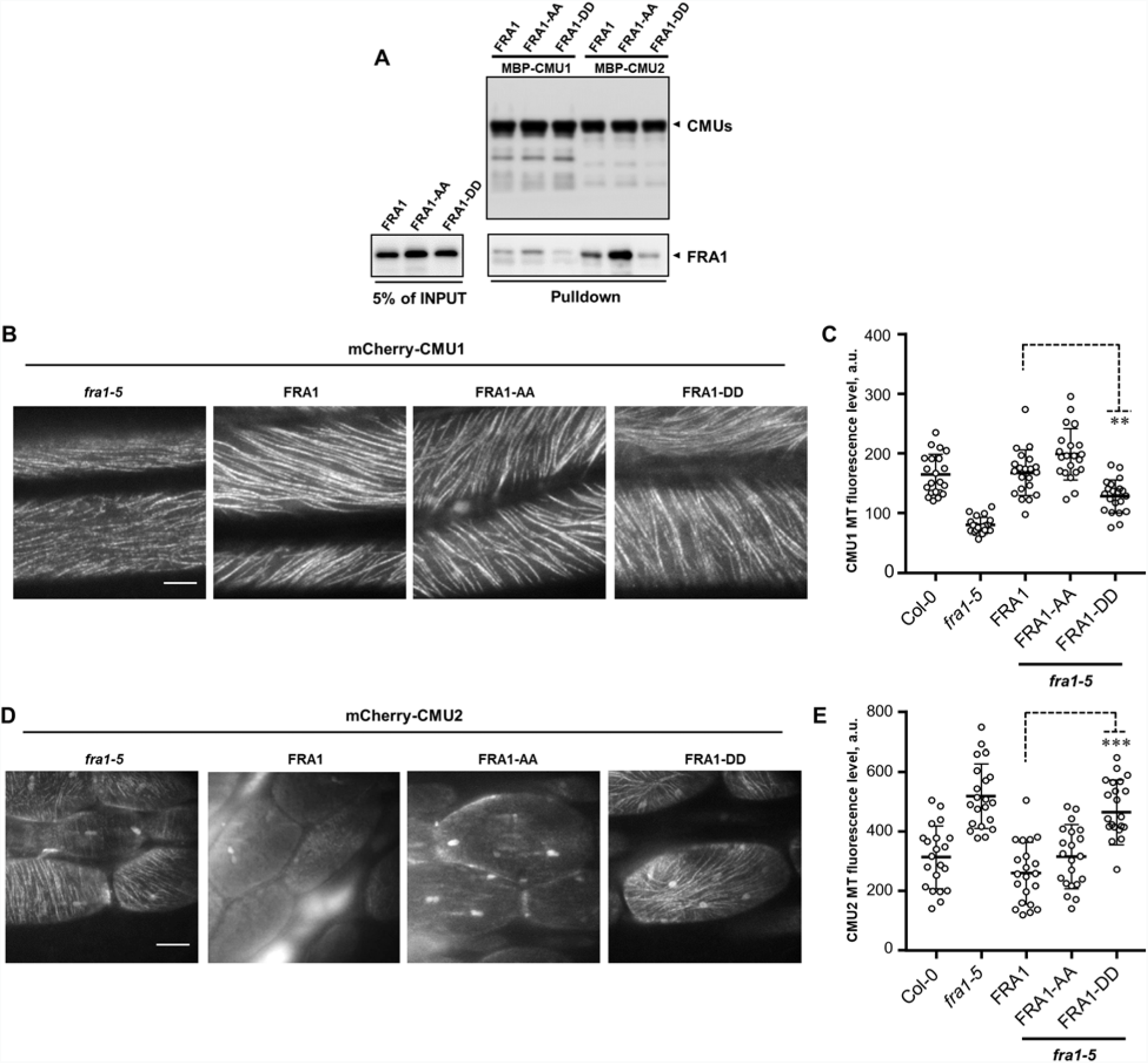
Phosphorylation of FRA1 regulates CMU1 and CMU2 localization on cortical microtubules. (A) Pulldown of MBP-CMU1 and MBP-CMU2 incubated with equal amounts of either wild-type FRA1-CT, FRA1-AA or FRA1-DD. Immunoblotting was performed with anti-MBP and anti-FRA1 antibodies. (B-E) Fluorescence micrographs (B and D) and quantification (C and E) of mCherry-CMU1 and mCherry-CMU2 signal on cortical microtubules in hypocotyl epidermal cells of 4-d old seedlings. Values are mean ± S.D. (3-4 microtubules were quantified per cell, total 8 cells from 3 seedlings). Scale bar = 10 µm for (B) and 5 µm for (D). Asterisks indicate significant difference between the indicated genotypes as determined by ANOVA, **p < 0.001, ***p < 0.0001.

## DISCUSSION

Cell wall assembly poses a systems engineering problem to cells. One of the key challenges is to coordinate the deposition of cellulose and matrix polysaccharides since they are synthesized at different cellular locations. This logistics challenge becomes more acute during periods of rapid growth when cells boost the abundance and activity of both CSCs and FRA1 kinesin to meet the increased demand for cellulose and matrix polysaccharides, respectively (Crowell et al., 2009; Zhu et al., 2015; Ganguly et al., 2018). An increase in the number of motile CSCs in the plasma membrane could be deleterious because they would predictably deflect greater numbers of cortical microtubules and thus potentially disrupt array organization. Therefore, increased cellulose synthesis needs to be accompanied by increased positional stability of cortical microtubules. Here, we show that the FRA1 kinesin physically interacts with CMU proteins and promotes binding of CMU1 to cortical microtubules. As a result, an increase in the amount of FRA1 in growing cells would simultaneously enhance the resistance of cortical microtubules to lateral displacement by CSCs (Figure 9A). The transport activity of FRA1 combined with its ability to bolster the lateral stability of cortical microtubules through CMUs provides a mechanism to coordinate the oriented deposition of cellulose and matrix polysaccharides.

**Figure 9.**
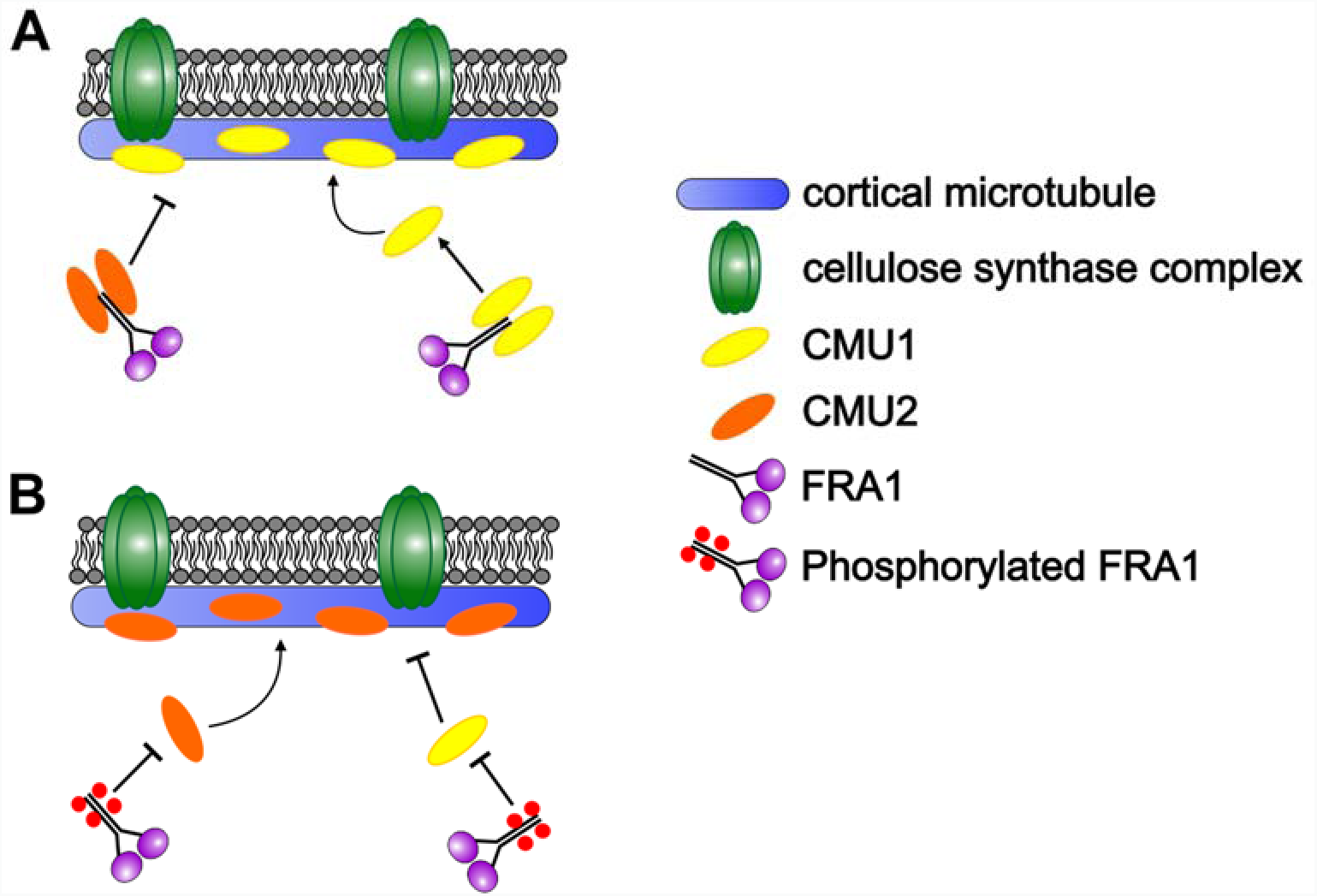
Model of how regulated interaction of FRA1 with CMU1 and CMU2 contributes to the lateral stability of cortical microtubules. (A) In seedlings, interaction with FRA1 promotes CMU1 binding to cortical microtubules but inhibits CMU2 binding to cortical microtubules. An increase in FRA1 levels would thus lead to more CMU1 binding and greater lateral stability of cortical microtubules. (B) In adult plants, the tail region of FRA1 is proposed to be phosphorylated by CKL6, which inhibits its interaction with both CMU1 and CMU2. This promotes cortical microtubule localization of CMU2 but inhibits CMU1 from doing so. Hence, phosphorylation provides a mechanism to control the relative amount of CMU1 and CMU2 on cortical microtubules.

CMUs resemble mammalian kinesin light chains (KLCs) because they are predicted to contain N-terminal heptad repeats and C-terminal tetratricopeptide repeats (Burstenbinder et al., 2013). In mammals, KLCs interact directly with the tail domain of kinesin-1 and modulate motor activity (Verhey et al., 1998; Cai et al., 2007; Wong and Rice, 2010). In addition, KLCs are also implicated as cargo adaptors for kinesin-1 (Stenoien and Brady, 1997; Khodjakov et al., 1998; Gyoeva et al., 2000; Morfini et al., 2002). We found that CMUs too directly interact with the tail domain of FRA1, however they do not regulate the motility of FRA1. CMUs are also unlikely to couple cargo to motile FRA1 molecules because CMUs appear as static puncta on cortical microtubules (Liu et al., 2016). Given these differences, CMUs are unlikely to function as plant counterparts of KLCs.

CMUs are microtubule-associated proteins (MAPs) by virtue of their microtubule localization *in vivo* and direct microtubule binding *in vitro* (Liu et al., 2016). Accruing evidence suggests that certain MAPs serve as positive or negative regulators of kinesins. For example, tau and MAP2 proteins both inhibit kinesin-1 motility (Heins et al., 1991; Seitz et al., 2002; Vershinin et al., 2007; Dixit et al., 2008). In contrast, MAP7 recruits kinesin-1 to microtubules and activates motility (Barlan et al., 2013; Tymanskyj et al., 2018; Chaudhary et al., 2019; Hooikaas et al., 2019). Similarly, the MAP65 family of microtubule bundling proteins recruit kinesin-4 to antiparallel microtubule overlaps in both plants and animals (Bieling et al., 2010; de Keijzer et al., 2017). However, we found that CMUs do not influence the amount, microtubule localization or motility of the FRA1 kinesin during interphase. Instead, we found that FRA1 regulates microtubule localization of CMUs, providing an example of a motor protein that modifies the behavior of its track. This activity appears to require direct binding of CMUs to FRA1 because inhibitory phosphomimetic mutations in the tail domain of FRA1 significantly attenuated FRA1’s impact on microtubule localization of CMUs. How FRA1 affects the microtubule binding of CMUs remains to be studied. One possibility is that FRA1 directly regulates CMU’s microtubule binding through an allosteric effect. Alternatively, FRA1 might indirectly achieve this outcome by influencing the binding of CMUs to other regulatory proteins. A plant-specific IQ67 domain (IQD) containing protein called IQD1 has been shown to bind to CMUs and regulate their microtubule localization (Burstenbinder et al., 2013). Therefore, it will be interesting to determine whether FRA1 affects the binding of CMUs and IQD1.

We found that roots of the *cmu1* mutant grow in a skewed manner, in agreement with the previously reported cell file twisting phenotype in hypocotyls of the *cmu1* mutant (Liu et al., 2016). In contrast, the *cmu2* seedlings did not show any measurable growth defects, at least under normal growth conditions. The *cmu1cmu2cmu3* triple mutant seedlings were also indistinguishable from wild-type control with the exception of having skewed roots similar to the *cmu1* single mutant. Together, these data indicate that CMU1 plays the primary role in axial growth of seedlings. None of the *cmu* mutant combinations showed significant growth defects in adult plants. However, when introduced into the *fra1-5* background, the *cmu* lesions exacerbated the growth and fertility defects of the *fra1-5* mutant. Thus, *fra1-5* provides a sensitized genetic background to study the contribution of cortical microtubule lateral stability conferred by CMUs to the growth and fecundity of plants.

Our genetic experiments revealed that *CMU1* and *CMU2* contribute additively to stem growth. We found that *CMU3* does not contribute to this phenotype, probably because CMU3 protein is undetectable in plants. Reciprocal pollination experiments demonstrated that reduced fertility of the *fra1-5* and *fra1-5cmu1cmu2* mutants was due to defective pistil and pollen functions. Successful pollen-pistil interactions involve pollen adhesion, hydration and germination on the stigma surface to produce a pollen tube which then penetrates the stigma and elongates into the transmitting tract of the style on its way to the ovary (Kandasamy et al., 1994). These steps require secretion of specific degradative enzymes, pollen tube growth promoting molecules and guidance factors into the cell wall of pistil tissues (Hiscock et al., 1994; Cheung et al., 1995; Lavithis and Bhalla, 1995; Hiscock et al., 2002; Okuda et al., 2009; Samuel et al., 2009). The reduced receptivity of the *fra1-5* and *fra1-5cmu1cmu2* pistils might be due to compromised secretion of these factors. It is also possible that cell walls of the transmitting tract of *fra1-5* and *fra1-5cmu1cmu2* pistils have altered composition and/or structure which might impede pollen tube growth. On the pollen side, our results suggest that FRA1 contributes to pollen tube elongation, perhaps by facilitating secretion of wall material at the growing tip. Furthermore, pollen viability was significantly reduced in the *fra1-5cmu1cmu2* mutant, indicating that proper cell wall production is important for pollen development.

Comparison of the cortical microtubule signal of mCherry-CMU1 and mCherry-CMU2 in wild-type and *fra1-5* hypocotyl epidermal cells revealed that FRA1 promotes CMU1 binding to cortical microtubules but inhibits CMU2 binding to cortical microtubules. As a result, wild-type seedlings have much higher levels of CMU1 on cortical microtubules compared to CMU2. The striking reduction in microtubule-associated CMU1 in the *fra1-5* mutant is likely the reason for the increased lateral deflections of cortical microtubules in *fra1-5* than in wild-type. In addition, even though both *CMU1* and *CMU2* are expressed in seedlings, only the *cmu1* mutant showed substantial skewing of the primary root, probably because FRA1 inhibits the binding of CMU2 to cortical microtubules. Conversely, an increase in the amount of CMU2 on cortical microtubules in the *fra1-5* mutant might partially compensate for the lack of CMU1, providing a plausible explanation for partial suppression of the root skewing phenotype in the *fra1-5cmu1* double mutant compared to the *cmu1* single mutant.

Arabidopsis protein phosphorylation datasets indicated that the tail region of FRA1 is phosphorylated at residues Thr687 and Thr694. Here, we show that CKL6 physically binds to the FRA1 tail region and can mediate this phosphorylation. Mutating Thr687 and Thr694 to alanine did not eliminate phosphorylation of the FRA1 tail region by CKL6, indicating that there are additional residues that serve as substrates for CKL6. In addition, we cannot exclude the possibility that other kinases phosphorylate FRA1 at these or other tail domain locations *in vivo*. Notably, phosphorylation of the FRA1 tail region did not affect its abundance or motility. Therefore, these phosphorylation events are not working to regulate motor activity. Rather, they regulate the binding of CMUs to FRA1 and consequently the extent of CMU localization to cortical microtubules. Since interaction with FRA1 differentially affected the cortical microtubule localization of CMU1 and CMU2, phosphorylation of the FRA1 tail region provides a mechanism to alter the balance of CMU1 and CMU2 along cortical microtubules. We found that while CMU1 is the major contributor to seedling growth, CMU2 is important for inflorescence stem growth in adult plants. The switch between CMU1 and CMU2 during seedling to adult plant transition could be achieved by phosphorylation of the FRA1 tail region (Figure 9B). Regulated interaction between FRA1 and CMUs may also be important for the development and function of plant reproductive tissues. Taken together, our data provide new insight into how mechanisms for the oriented deposition of cellulose and matrix polysaccharides functionally intersect at the cortical microtubule interface.

## METHODS

### Plant Material and Growth

*Arabidopsis thaliana* L. (Heynh), Columbia accession, was used throughout. The *cmu1* and *cmu2* mutants were isolated from T-DNA insertion lines, SAIL_335_B08 and SALK_148296C, respectively, obtained from ABRC (Alonso et al., 2003). Homozygous mutants were identified using primers listed in Supplemental Table 1. The *fra1-5* mutant was described in Zhu et al., 2015. For growth on plates, seeds were sterilized with 5% (v/v) bleach for 10 min, rinsed with sterile water four times and planted on 0.5X Murashige and Skoog medium (Caisson Laboratories). Seeds were stratified at 4 °C for 2 d and then germinated at 20 °C with 16 h of 120-140 µmol light. Live imaging used 4-d-old light-grown seedlings unless otherwise stated. Root length and root skewing angle were measured using the Fiji ImageJ package (Schindelin et al., 2012) on 5-d-old light-grown seedlings. For growth in soil, seeds were grown under continuous light at 120-140 µmol intensity, 70% humidity and 21°C after stratification at 4 °C for 2 d. Stem height was measured with a ruler.

### RT-PCR Analysis

Total RNA was extracted from basal internodes of 4-week-old plants using Omega Plant RNA Mini Kit (Omega Bio-Tek). About 500 ng DNAse treated RNA was used for cDNA synthesis using qScript cDNA supermix (Quanta BioSciences). Primers for RT-PCR were designed using sequence downstream of the T-DNA insertion sites and are listed in Supplemental Table 1.

### Generation of CMU1 and CMU2 Constructs and Transgenic *Arabidopsis* Plants

Primers used for construct assembly are listed in Supplemental Table 1. The *pCMU1::mCherry-CMU1 and pCMU2::mCherry-CMU2* constructs were generated using about 2kb *CMU1* or *CMU2* promoter, *mCherry* cDNA and full-length genomic DNA of *CMU1* or *CMU2* genes, respectively. The constructs included a 36-bp linker sequence between *mCherry* and *CMU1* or *CMU2.* The assembled constructs were ligated into pCAMBIA 3300 vector and introduced into Arabidopsis plants expressing the GFP-TUB6 microtubule marker (gift from Prof. Hashimoto) via Agrobacterium-mediated floral dip transformation. Transgenic plants were selected using 10 µg/ml phosphinothricin and T2 lines expressing single copy of the transgene were used for imaging. The *pFRA1::FRA1-tdTOMATO* construct has been described in Zhu et al., 2015. The *pFRA1::FRA1-EGFP* construct was made by replacing the 3X-GFP in the *pFRA1::FRA1-3X-GFP* (Zhu et al., 2015) by a single *EGFP* gene. The T687A-T694A and T687D-T694D mutations were constructed by megaprimer method using primers listed in Supplemental Table 1.

### Live Imaging of Arabidopsis Seedlings

*Arabidopsis* seedlings were imaged using variable-angle epifluorescence microscopy (Konopka and Bednarek, 2008). Live imaging of mRuby-TUB6 decorated microtubules were conducted using variable-angle epifluorescence microscopy. Four-day-old seedlings were mounted in 0.5X Murashige and Skoog media between two layers of double-sided adhesive tape. Unless otherwise stated, epidermal cells in the apical or sub-apical region of the hypocotyl were imaged. mRuby was excited using 5 mW 561-nm diode-pumped solid-state laser (Melles Griot). To image FRA1-EGFP and FRA1-tdTomato, specimens were excited using 3 mW 488-nm and 5 mW 561-nm diode-pumped solid-state lasers (Melles Griot), respectively. Images were collected using 100X (NA 1.45) objective and back-illuminated electron-multiplying CCD camera (Hamamatsu, ImageEM) at 1-s for single-color imaging and 3-s for two-color imaging. For drug treatments, 4-day-old light-grown seedlings were incubated with either 20 µM oryzalin (Supelco Analytical) for 3 h or 2 µM latrunculin B (Enzo Life Science) for 2 h before imaging.

### Yeast Two-Hybrid Assay

The initial yeast two-hybrid screen that identified CMU1 and CMU2 as potential interaction partners of FRA1 was conducted by Hybrigenics Services. Directed yeast two-hybrid experiments were performed according to the ProQuest™ Two-Hybrid System with Gateway® Technology manual (Invitrogen). Full-length CMU1 and CMU2 along with a FRA1 C-terminal fragment (amino acids 616-1035) were first cloned into the entry vector pENTR 11 with enzymes Kpn1 and Not1 for CMU1 and CMU2 and Kpn1 and EcoR1 for FRA1 (for primers, see Supplemental Table 1). From the entry clones, CMU1, CMU2 and FRA1-tail domain were then cloned into pDEST32 and pDEST22 vectors, respectively, using LR Clonase II enzyme (Invitrogen). These constructs together with all the controls were transformed into the MaV203 Competent Yeast Cells (*Saccharomyces cerevisiae*) and selected on SD-Leu-Trp plates. For each set, at least four independent colonies were used for yeast two-hybrid assay on SD-Leu-Trp-His plates with and without 3-amino-1,2,4-triazole (Sigma). Colonies were imaged every 24 h post plating for 4-5 days.

### *In Vitro* Pull-Down Experiments

CMU1, CMU2 and CMU3 were cloned into pMAL:c5x vector to obtain N-terminal MBP-tagged fusion proteins. The FRA1 C-terminal tail fragment (amino acids 616-1035) was cloned into a pTEV vector for a C-terminal 6xHis fusion construct. Plasmids were transformed into BL21-DE3-RIPL competent cells (Agilent) and induced with 0.5 mM IPTG for 4 hours. Cells were lysed with Thermo Scientific Pierce B-PER Bacterial Protein Extraction Reagents (1 ml B-Per, 2 µl DNase1, 3 µl Lysozyme, 1X protease inhibitor, 1 mM PMSF). For the pull-down experiments, 40 µl of MBP-Trap agarose beads (Chromotek) were first incubated with either MBP, MBP-CMU1, MBP-CMU2 or MBP-CMU3 (~5 ug each) at 4°C on a shaker (speed 10 rpm) for 4-6 h to allow binding. Then, purified FRA1 tail fragment was added (~5 µg) into each tube and incubated for an additional 10-12 h as before. Subsequently, the beads were washed at least 5 times with 1X PBS + 0.1% (v/v/) Tween-20. Bound proteins were isolated from the beads by adding 1X SDS loading dye, separated by SDS-PAGE and transferred to 0.45 µm PVDF membrane (Thermo Scientific). The blots were probed with a monoclonal anti-MBP antibody from DSHB, Iowa State University, DSHB-MBP-2A1 (1:5,000), and a polyclonal anti-FRA1 antibody (1:2,000) described previously (Zhu et al., 2015). For secondary antibodies, anti-rabbit IgG HRP (1:10,000, Jackson Immuno Research) and anti-mouse IgG HRP (1:10,000, Jackson Immuno Research) were used. Detection was conducted using SuperSignal West Dura chemiluminescence substrate (Thermo Scientific). CKL3 and CKL6 were cloned into the pMAL:c5x vectors to obtain N-terminal MBP-tagged fusion proteins. Pull-down experiments with 6xHis-tagged FRA1 tail fragment were performed using the above protocol.

### *In Vivo* Pull-Down Experiments

Native promoter-driven *CMU1* and *CMU2* constructs were introduced into the pCAMBIA3300 vector using primers listed in Supplemental Table 1. The constructs were transformed into Col-0 plants and homozygous transgenic plants were selected. For the pull-down assay, 7-day-old seedlings were homogenized in a mortar using lysis buffer (20 mM HEPES pH 7.5, 40 mM KCl, 1 mM EDTA, 0.1% (v/v) Triton-X 100, 1X protease inhibitor cocktail, 1 mM PMSF, 10% (v/v) glycerol). The homogenate was then centrifuged twice at 10,000 rpm for 10 min and incubated with 20 ul of anti-mCherry Trap agarose beads (Chromotek) for 12-16 h at 4°C on a rocking shaker. Beads were then washed at least 5 times with 1X PBS + 0.1% (v/v) Tween-20 solution. Bound proteins were isolated from the beads by adding 1X SDS loading dye, separated by SDS-PAGE and detected by immunoblotting using anti-mCherry and FRA1 antibody.

### Pollen Viability Tests

Pollen from 6 fully open flowers of 4-5 week old plants were incubated in 50 µl pollen germination medium (10% sucrose, 0.01 mM Boric Acid, 5 mM CaCl_2_, 5 mM KCl, 1 mM MgSO_4_, pH 7.5) on a slide for 10 min and then stained with 1 µg/ml propidium iodide for 15 min. Stained pollen were imaged using a UV-fluorescence microscope with RFP filter and 20X lens.

### Pollination and Aniline Blue Staining

Pistils from stage 12 flowers were exposed by removing sepals, petals and stamens and kept upright in 1% (w/v) agar plates. Any pistils with pollen already on them were discarded. Virgin pistils were pollinated using anthers from fully open flowers. After 2-h at room temperature, pistils were fixed using acetic acid:ethanol (3:1 v/v) at 60 °C for 5 minutes. The fixative solution was replaced with 1N NaOH and incubated at 60°C for 15 minutes to soften the tissue. After rinsing with water, the samples were stained with 0.1% decolorized aniline blue, which stains callose. Stained pistils were imaged under a UV-fluorescence microscope with a DAPI filter. Pollen tube length was measured from the stigma tip to where the majority of pollen tubes ended within the style using the Fiji ImageJ package (Schindelin et al., 2012). For each pistil, the pollen tube lengths were measured three times and the average was used.

### *In Vitro* Kinase Assay

The FRA1 tail fragment (amino acids 616-1035) was cloned into the pMAL:c5x vector (MBP tag) with Sal1 and EcoR1 enzymes and CKL3 and CKL6 were cloned in the pTEV vector (6xHIS tag) with Nde1 and BamH1 sites. For the kinase assay, MBP, MBP-FRA1-tail, and MBP-FRA1(AA)-tail were used as potential substrates for CKL3-6xHIS and CKL6-6xHIS. For each assay, 2µg of substrate and 500 ng of kinase were incubated in 1X Kinase Buffer (25 mM Tris-HCl pH 7.5, 2 mM DTT, 50 µM ATP, 12 mM MgCl_2_) supplemented with 50 µM ATP and 1X phosphatase inhibitor (EMD Biosciences, phosphatase inhibitor complex set III, Cat# 524627) in a 50 µl reaction. Reactions were carried out at 30°C for 45 min and then terminated by adding 4X SDS-Page loading buffer and heated at 75 °C for 6-8 min. Bands were separated by SDS-PAGE and detected via immunoblotting using anti-phospho serine/threonine antibody (ECM Biosciences, Rabbit polyclonal Anti-phosphoserine-threonine antibody, Cat# PP2551) at 1:2500 dilution and anti-rabbit goat secondary antibody with HRP, at 1: 10,000 dilution.

## AUTHOR CONTRIBUTIONS

AG, CZ and RD designed the research and analyzed the data. AG, CZ and WC performed the research. AG and RD wrote the article. All authors read and approved the final version of the manuscript.

## ACKNOWLEDGEMENTS

We thank Graham Burkart for the mRuby-TUB6 microtubule marker line used in this study; Katharina Bürstenbinder for sharing unpublished data on FRA1-CMU interaction and Yanbing Wang for help with pollination experiments. This work was supported by NSF grant 1453726 to R.D.

