## Supplemental figures and table for "FRA1 modulates cortical microtubule localization of CMU proteins"

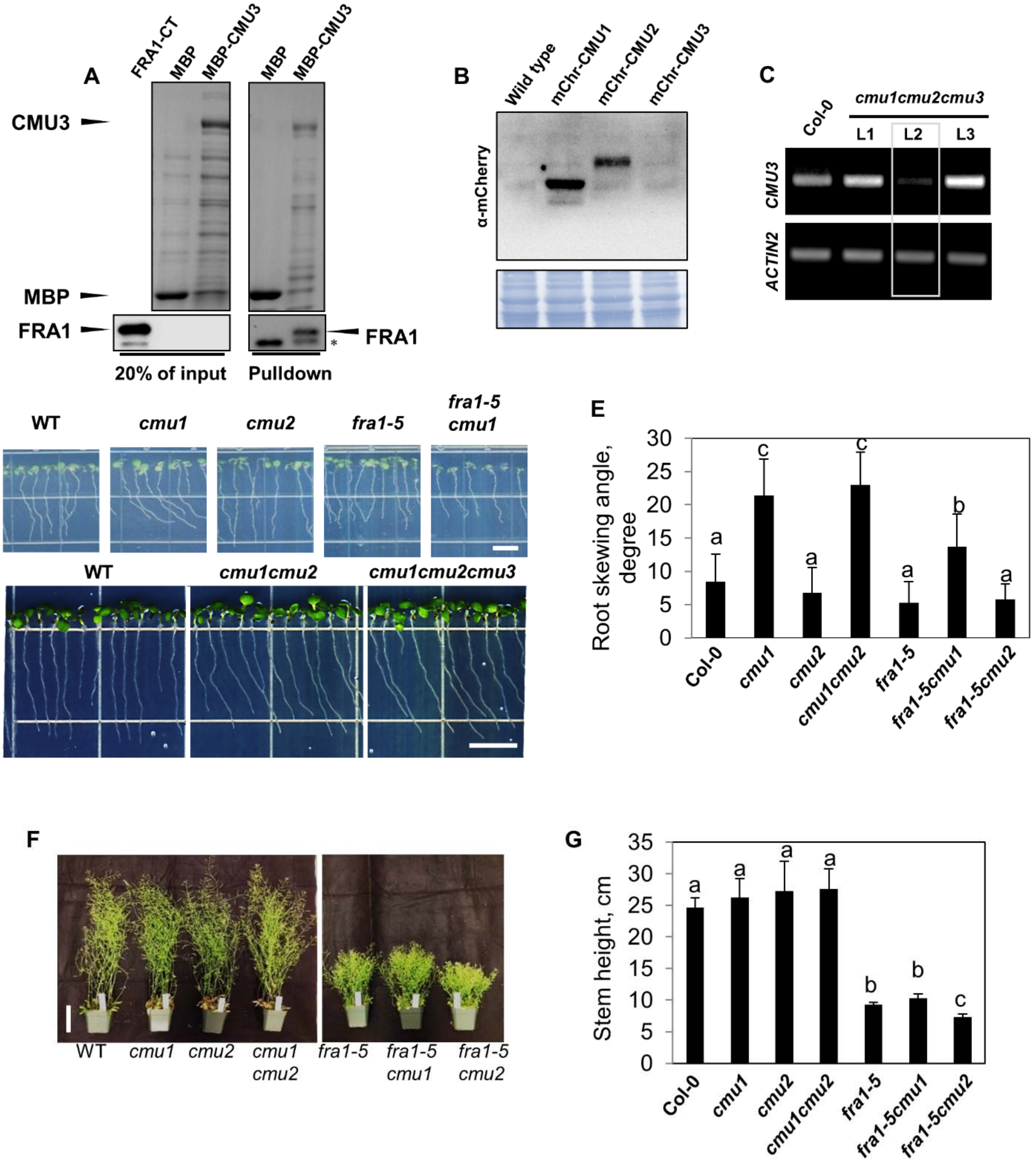


**Supplemental Figure 1.** Seedling and adult plant phenotypes of *cmu* and *fra1-5* mutant combinations.

(A) Pulldown of either MBP alone or MBP-tagged CMU3 incubated with equal amounts of FRA1-CT. Immunoblotting was performed with anti-MBP and anti-FRA1 antibodies. *denotes a non-specific band.

(B) Immunoblot of total protein extracts prepared from Col-0 plants expressing either mCherry-CMU1 (mChr-CMU1), mCherry-CMU2 (mChr-CMU2) or mCherry-CMU3 (mChr-CMU3) probed with anti-mCherry antibody. Coomassie-stained gel is shown below as a loading control.
(C) RT-PCR analysis of *CMU3* and *ACTIN2* expression in Col-0 and three independent *cmu1cmu2cmu3* transgenic lines. *ACTIN2* expression is used as a loading control. Line 2 (L2) has significantly reduced *CMU3* expression and was chosen for further analysis.
(D, E) Root images and quantification of root skewing angle of 5-day old light-grown seedlings. Values are mean ± SD from 20 seedlings for each genotype. Different letters on bars indicate significant difference as determined by ANOVA, $p < 0.05$. Scale bar = 0.5 cm.
(F, G) Whole-plant appearance and quantification of stem heights of plants 42 days after germination. Values are mean ± S.D. (n = 12 plants each). Different letters on bars indicate significant difference as determined by ANOVA, $p < 0.05$. Scale bar = 5 cm.

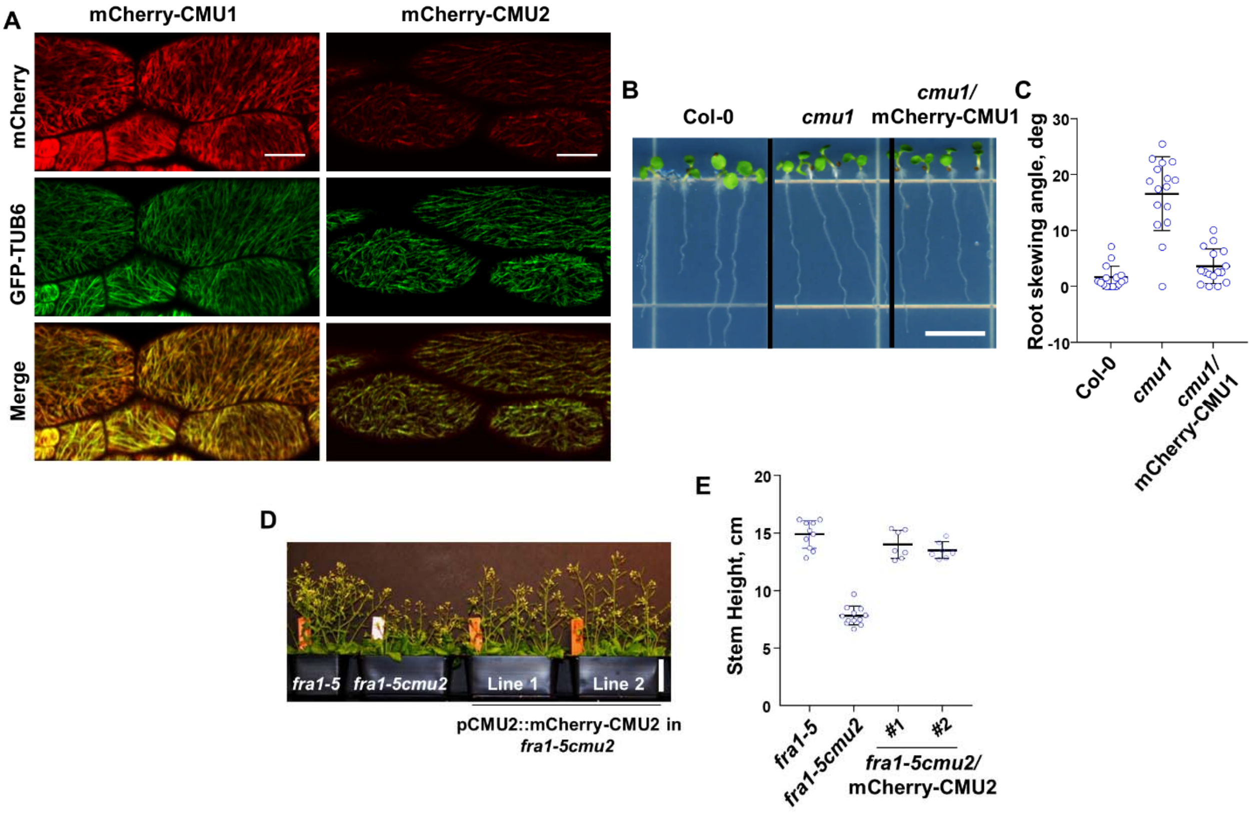


**Supplemental Figure 2**. Cortical microtubule localization of functional mCherry-labeled CMUs. (A) Fluorescence micrographs of hypocotyl epidermal cells from a 4-d old *Arabidopsis* seedlings expressing GFP-TUB6 and either mCherry-CMU1 or mCherry-CMU2. Scale bar = 5 µm.

(B, C) Root images and quantification of root skewing angle of 5-day old light-grown seedlings. Values are mean ± SD from >15 seedlings for each genotype. Scale bar = 0.5 cm

(D, E) Whole-plant appearance and quantification of stem heights of plants 56 days after germination. Values are mean ± S.D. (n = 12 plants each). Scale bar = 5 cm.

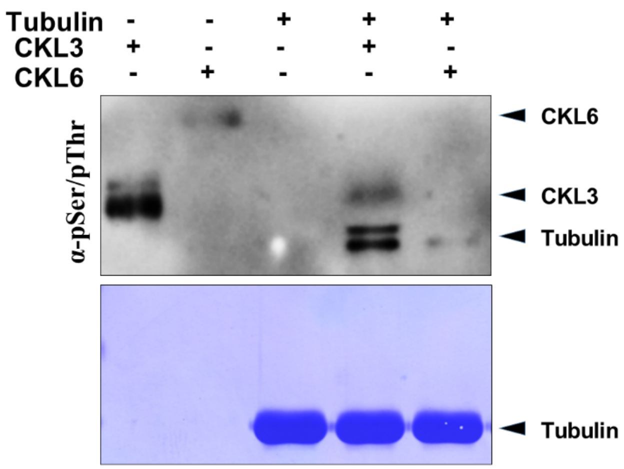


**Supplemental Figure 3**. CKL3 and CKL6 are functional kinases.
*In vitro* phosphorylation experiment with porcine tubulin incubated with either CKL3 or CKL6. Phosphorylation was detected by an anti-phospho serine/threonine antibody. Coomassie-stained gel is shown below as a loading control. CKL3 and CKL6 phosphorylate tubulin and show autophosphorylation.

**Supplemental Table 1**: Primers used in this study

| Gene ID | Gene name | Primer Sequence (5'-3') | Application |
|---|---|---|---|
| At5g47820 | FRA1 | GCTGCAGGTACAAATTGGATCAC | Mutant screening |
| | | TTTCTCGAATCTGTATCGGTGGC | |
| At4g10840 | CMU1 | TTGAGGATAAGCCAGGAC AG | |
| | | TCCTCTCTCAGCTTCAGAAC | |
| At3g27960 | CMU2 | TATAACTAGTATGGACGTAGGAGAGAGCAATG | |
| | | CCATACATGTCACCAAGCTGC | |
| At3G18780 | ACTIN2 | TCAATCATGAAGTGTGATGTGG | RT-PCR |
| | | AACGACCTTAATCTTCATGCTGC | |
| At4g10840 | CMU1 | GCAGGTTCTGAAGCTGAG AGAG | |
| | | CTTCGGAGGTCTTGCATTTGG | |
| At3g27960 | CMU2 | GAAGCTCAAATGGGTGTG GTTAC | |
| | | CAGCTTCATTGATCGCGTAACG | |
| At4g10840 | pCMU1 | TATAGGATCCAATCTTACGGATGGACTTCTATGC | Cloning in pCAMBIA1300 vector |
| | | TATACCCGGGGAATGTGTCTCTCTGTGGGAAG | |
| At3g27960 | pCMU2 | TATAGGATCCGATATGGGATTCGAGCGGC | |
| | | TATACCCGGGGGCCTCCAAAACTCACAACTC | |
| At4g10840 | CMU1 | TATAACTAGTATGCCAGCAATGCCAGGTCT | |
| | | TATAGAGCTCTCAGAACTTGAAACCGAGGC | |
| At3g27960 | CMU2 | TATAACTAGTATGGACGTAGGAGAGAGCAATG | |
| | | TATAGAGCTCTCAATAAACCGGTCTCTGTCC | |
| At1g27500 | CMU3 | TATAACTAGTATGATGGAAGGAGGGTCTGTTAATG | |
| | | TATAGAGCTCTTAACGAAGAGCTGAAGAAGAAGT | |
| At1g27501 | pCMU3 | TATAGGATCCGCAAGGAAAAATCGCTGACT | |
| | | TATACCCGGGTTTCTTTATTTCGAATAAACAGCTATCTG | |
| At4g10840 | CMU1 | TATAGCGGCCGCATGCCAGCAATGCCAGGTCT | pMAL:C5X (MBP Tag) expression vector |
| | | TATAGAATTCTCAGAACTTGAAACCGAGGC | |
| At3g27960 | CMU2 | TATAGCGGCCGCATGGACGTAGGAGAGAGCAATG | |
| | | TATAGAATTCTCAATAAACCGGTCTCTGTCC | |
| At1g27500 | CMU3 | TATAGCGGCCGCATGGAAGGAGGGTCTGTTAATG | |
| | | TATAGAATTCTTAACGAAGAGCTGAAGAAGAAGTGAG | |
| At4g10840 | CMU1 | TATAGGTACCGAATGCCAGCAATGCCAGGTCTCGTC | Yeast two-hybrid cloning |
| | | TATAGCGGCCGCTCAGAACTTGAAACCGAGGCTGG | |
| At3g27960 | CMU2 | TATAGGTACCGAATGGACGTAGGAGAGAGCAATGAG | |
| | | TATAGCGGCCGCTCAATAAACCGGTCTCTGTCC | |
| At1g27500 | CMU3 | GATTGAATGCATGCTAAACCCCTTCTCTCTTTTGTATTCC | amiRNA for CMU3 |
| | | GAAGGGGTTTAGCATGCATTCAATCAAAGAGAATCAATGA | |
| | | GAAGAGGTTTAGCATCCATTCATTCACAGGTCGTGATATG | |
| | | GAATGAATGGATGCTAAACCTCTTCTACATATATATTCCT | |
| | mCherry | TATACCCGGGATGGTGAGCAAGGGCGA | |

| | | | |
|---|---|---|---|
| | | TATAACTAGTTGCACCCGTAGTCGCAGCGGCTCCGGTTGTTGCGGCCTTGTACAGCTCGTCCATGC | Cloning in pCAMBIA1300 vector |
| | EGFP | TATAACGCGTTGCACCCGTAGTCGCAGCGGCTCCGGTTGTTGCGGCATGGTGAGCAAGGGCGAGG | |
| | | TATAGAGCTCTTACTTGTACAGCTCGTCC | |
| At5g47820 | FRA1 | CTCTTTCAACCTTTTGGCAGCCATTGCAGCCTCTTCTGCCTTCCTCTGAAGAAC | T687A-T694A |
| | | CTCTTTCAACCTTTTGTCAGCCATTGCAGCCTCTTCATCCTTCCTCTGAAGAAC | T687D-T694D |
| At1g27500 | CMU3 | GAGGAAAAGCTCGGGACAG | RT-PCR |
| | | CAAATCTGTGAGAGCCTGTTAA | |
| At4g28880 | CKL3 | TATACATATGGAACGCATCATCGGCGGCAAGTACAAGC | pMAL:C5X (MBP Tag) and pTEV (6X-HIS tag) expression vector |
| | | TATAGGATCCTTTCTCTGGAGACATGAATGACGAAGTCC | |
| At4g28540 | CKL6 | TATACATATGGACTTGAAAATGGATAATGTTATTGGG | |
| | | TATAGGATCCTTTGCGGATCGAAAGAAGCTCGAAGC | |
| At5g47820 | FRA1-CT | TATAGTCGACGCTCGGAGGTTACAAGATGA | pMAL:C5X (MBP Tag) expression vector |
| | | TATAGAATTCTTACATGATCTTATTAGGTAGA | |
| SAIL line | Left Border | GAATTTCATAACCAATCTCGA TACAC | Mutant screening |
| SALK line | Left Border | TGGTTCACGTAGTGGGCCATCG | Mutant screening |
